# Computational and experimental insights into the chemosensory navigation of *Aedes aegypti* mosquito larvae

**DOI:** 10.1101/585075

**Authors:** Eleanor K. Lutz, Tjinder S. Grewal, Jeffrey A. Riffell

## Abstract

Mosquitoes are prolific disease vectors that affect public health around the world. Although many studies have investigated search strategies used by host-seeking adult mosquitoes, little is known about larval search behavior. Larval behavior affects adult body size and fecundity, and thus the capacity of individual mosquitoes to find hosts and transmit disease. Understanding vector survival at all life stages is crucial for improving disease control. In this study we use experimental and computational methods to investigate the chemical ecology and search behavior of *Aedes aegypti* mosquito larvae. We show that larvae do not respond to several olfactory cues used by adult *Ae. aegypti* to assess larval habitat quality, but perceive microbial RNA as a potent foraging attractant. Second, we demonstrate that *Ae. aegypti* larvae use a strategy consistent with chemokinesis, rather than chemotaxis, to navigate chemical gradients. Using computational modeling, we further show that chemokinesis is more efficient than chemotaxis for avoiding repellents in ecologically relevant larval habitat sizes. Finally, we use experimental observations and computational analyses to demonstrate that larvae respond to starvation pressure by optimizing exploration behavior. Our results identify key characteristics of foraging behavior in a disease vector mosquito, including the identification of a surprising foraging attractant and an unusual behavioral mechanism for chemosensory preference. In addition to implications for better understanding and control of disease vectors, this work establishes mosquito larvae as a tractable model for chemosensory behavior and navigation.

## Introduction

The mosquito *Aedes aegypti* is a global vector of diseases such as Dengue, Zika, and Chikungunya [1]. This synanthropic mosquito is evolutionarily adapted to human dwellings, with some populations breeding exclusively indoors [2, 3]. The urban microhabitat is a fascinating environment with unique climatic regimes, photoperiod, and resource availability. In response to these selective pressures, successful synanthropic animals including cockroaches [4], rats [5], and crows [6] exhibit many behaviors absent in non-urbanized sibling species. Understanding these behaviors is of major importance to public health. Throughout human history, synanthropic disease vectors have caused devastating pandemics like the Black Death, which killed an estimated 30-40% of the Western European population [7, 8]. Like rats or cockroaches, adult *Ae. aegypti* mosquitoes exhibit many behavioral adaptations to human microhabitats [2, 9]. However, comparatively little is known about larval adaptations. The larval environment directly affects adult body size [10, 11], fecundity [11], and biting persistence [12], and understanding vector survival at all life stages is crucial for improving disease control [13]. Despite growing interest [14, 15, 16], it remains an open question of how environmental stimuli affect larval behavior to regulate these responses and processes.

In addition to the above public health implications, the behavior of synanthropic mosquito larvae is fascinating from a theoretical search strategy perspective. *Ae. aegypti* larvae are aquatic detritivores that live in constrained environments such as vases and tin cans [10]. In such limited environments, do larva exhibit a chemotactic search strategy (in which animals change their direction of motion in response to a chemical stimuli), or do they use a chemokinetic response (in which animals change a non-directional component of motion, such as speed or turn frequency, in response to a chemical stimuli) [17], or a purely stochastic behavior, akin to a random walk? Mechanistic understanding of larval foraging behavior may provide insight into chemosensory systems controlling the behavior as well as the evolutionary adaptations for these systems in synanthropic environments.

In this work, we investigate larval *Ae. aegypti* behavior from a chemical ecological and search theory perspective. First, we explore the chemosensory cues involved in larval foraging. Although many olfactory cues are used by adult females to select oviposition sites [18], it is unclear if larvae and adults use the same chemicals to assess larval habitat quality. Second, we consider larval search behavior in spatially restricted environments using empirical data and computational modeling. Our work identifies the functional loss of chemotaxis in foraging larvae - a fascinating example of how environmental restrictions can drive the evolution of animal behavior. We further identify microbial RNA as a potent and unusual larval foraging attractant.Together, our results identify *Ae. aegypti* larvae as an exciting outlier in biological search theory, and highlights the importance of investigating synanthropic disease vectors at all life history stages.

## Results

### Effects of sex, physiological state, and circadian timing on larval physiology

Behavioral experiments in insects have demonstrated the importance of circadian timing, starvation, and age [19]. However, little is known about the effects of these variables on *Ae. aegypti* larvae. To better understand the baseline characteristics of our study organism, we used machine vision to track individual *Ae. aegypti* larvae in a custom arena (Fig 1A) and investigated the effects of nutritional state and sex on baseline larval behavior recorded before each experiment. For both fed and starved animals, female larvae were larger than males (fed larvae: n=120♀, 128♂, p<0.0001; starved larvae: n=79♀, 89♂, p=0.008, Fig S1A). Starved larvae were also smaller than fed animals for both females (p<0.0001) and males (p=0.015, Fig S1A). Because adult *Ae. aegypti* exhibit crepuscular activity [10], we also investigated the effects of circadian timing on larval behavior. We found no effects of circadian timing on larval movement speed, time spent moving, or time spent next to arena walls - supporting previous findings that mosquito larvae, unlike adults, exhibit little behavioral variation during the day [20, 21] (p=1, p=1, p=1, Fig S1B-D).

**Figure 1:**
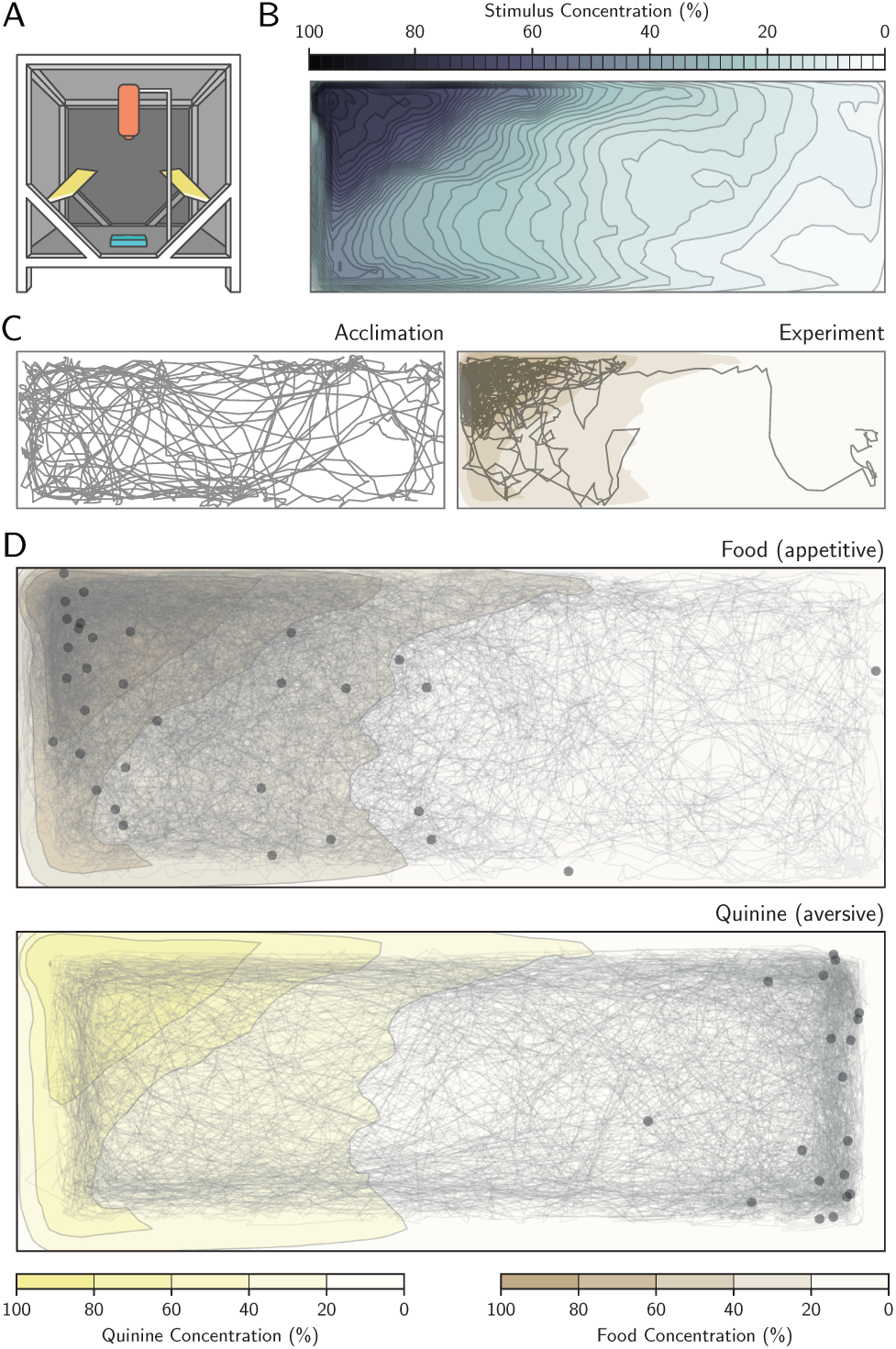
Quantifying the chemosensory environment in naturalistic larval habitat sizes. A: Diagram of experimental conditions, adapted from [22], including a Basler Scout Machine Vision GigE camera (orange), infrared lighting (yellow) and a behavior arena (blue). B: Chemosensory diffusion map of the behavior arena at the end of the 15 minute experiment. C: Example of an individual larval trajectory during the 15 minute acclimation phase (left). Trajectory of same individual during the 15 minute experiment phase, responding to food added to the left side of the arena (right). D: Trajectory of all starved animals presented with food (top) or quinine (bottom). Although trajectories are shown aggregated into one image, all animals were tested individually. Scatter points show the position of each animal at the end of the experiment.

### Quantifying the chemosensory environment in naturalistic larval habitat sizes

Previous research has shown that other species of mosquito larvae detect many different chemosensory stimuli [23]. In *Ae. aegypti* it is unclear what chemical signals, if any, larvae use to navigate their environment. Nevertheless, chemosensory cues may be essential in avoiding predation or foraging efficiently. Using our arena and machine vision methods, we investigated larval preference for six putatively attractive and aversive chemosensory cues. First, we experimentally verified the chemical diffusion in the arena and found that larval activity significantly influenced the distri-bution of stimuli within the arena (p<0.0001). We next created a chemical diffusion map for analyzing stimuli preference using only experiments containing larvae (Fig 1B, Fig S2A-D). For chemosensory stimuli, we used predicted attractive stimuli including a 0.5% mixture of food (Hikari Tropic First Bites fish food) suspended in water, as well as food extract filtered through a 0.2*μ*m filter to remove solid particulates. Quinine was used as a putative aversive stimulus (a bitter tastant aversive to many insects including *Drosophila melanogaster* and *Apis mellifera* [24, 25]). We also tested indole and o-cresol, two microbial compounds that attract adult mosquitoes for oviposition [26]. Finally, we examined the larval response to microbial RNA. RNA is required for *Ae. aegypti* larval survival [27], and nucleic acids attract larvae of several other mosquito species [28]. Moreover, dissolved RNA is released at high levels (*μ*g/h/L) from growing populations of microbes in freshwater habitats [29], and could provide valuable foraging information to omnivores such as *Ae*. *aegypti.* By contrast, other isolated macronutrients such as salts, sugars, and amino acids elicit little to no attraction [28].

### Physiological feeding state affects larval attraction towards ecologically relevant odors

For each of these seven stimuli, we compared the stimulus preference of larvae before and after stimulus addition (Fig 1C, Fig 2A). Preference was defined as the median concentration chosen by the larvae throughout the 15-minute experiment, normalized to behavior during the previous 15-minute acclimation phase. Starved larvae were attracted to food (n=32, p<0.0001) and spent significantly less time near the aversive cue quinine (n=19, p<0.0001). Food extract filtered through a 0.2*μ*m filter remained attractive (n=19, p=0.004), suggesting that larvae use small, waterborne chemical cues to forage. To further investigate these foraging cues, we next examined responses to microbial RNA, and found that RNA was significantly attractive (n=18, p=0.047). Addition of water - a negative control for mechanical disturbance - had no impact on larval positional preference (n=16, p=1). Although we expected indole and o-cresol, which are attractive to adult *Ae. aegypti*, to elicit attraction from larvae, neither odorant elicited a change in behavior from the acclimation phase (indole: n=20, p=1; cresol: n=25, p=1). Indole tested at a higher concentration (10mM) also had no effect (n=19, p=0.28). Together, these results suggest that larvae and adults may not nec-essarily rely on similar cues to assess larval habitat quality.

**Figure 2:**
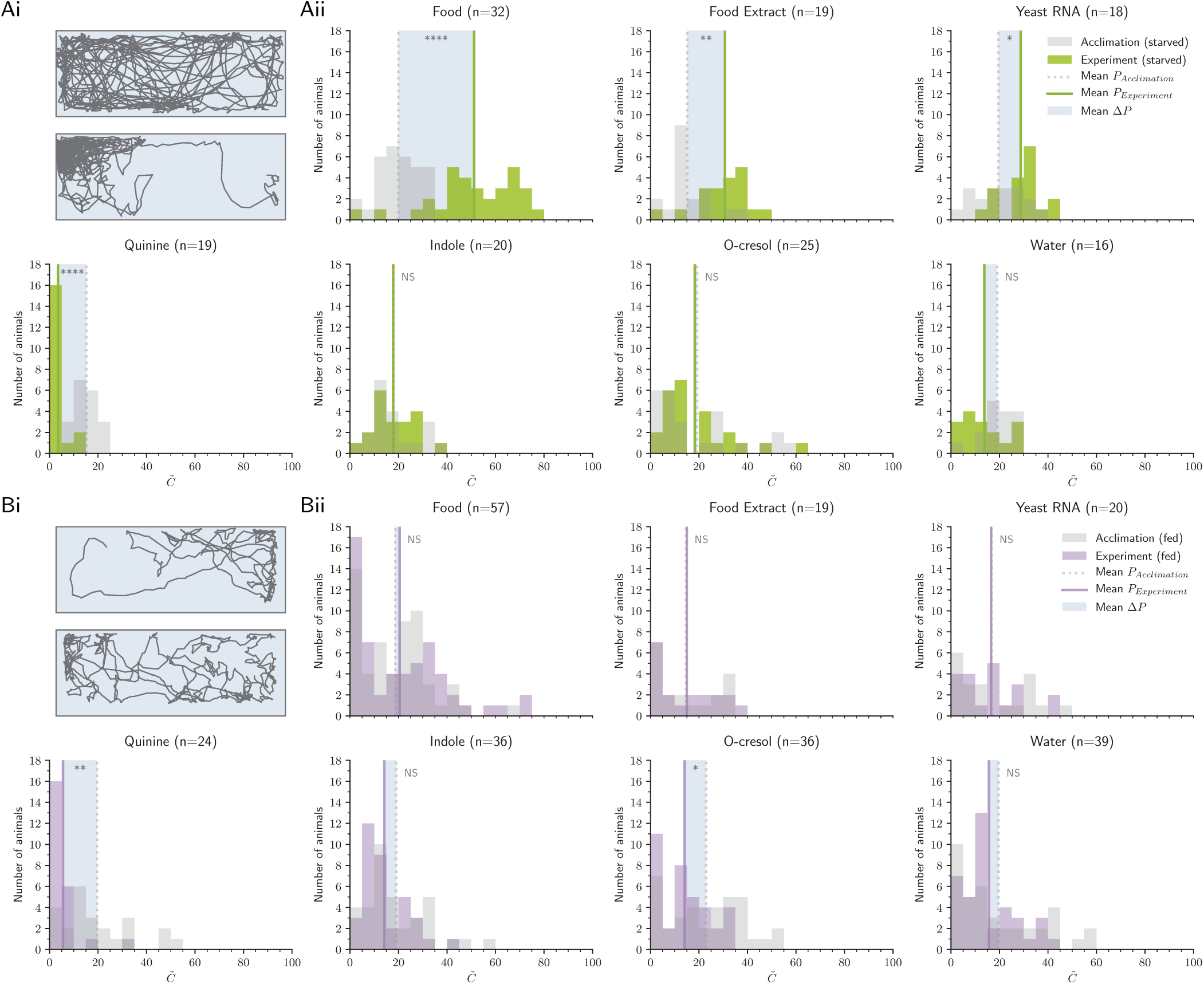
Physiological feeding state affects larval attraction towards ecologically relevant odors. Ai: Example trajectory of a starved larva during the acclimation (top) and the experiment phase (below), responding to food stimulus. Aii: Distribution of larvae during the acclimation phase (grey) and experiment phase (green), mean concentration 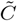. The shaded box visualizes the mean ΔP. Note that due to the unequal distribution of high and low concentration areas in the behavior arena, animals naturally appear to distribute near lower concentrations when no stimulus is present. Bi: Example trajectory of a fed larva during the acclimation (top) and experiment phase (below), responding to food stimulus. Bii: Distribution of fed larval preference during the acclimation (grey) and experiment phase (purple). In Aii and Bii, asterisks denote the significance level of paired-sample Welch’s t-tests comparing acclimation P and experiment P (NS: not significant). N values reported next to each stimulus describe the number of animals in the treatment.

The physiological feeding state of an adult mosquito has a strong impact on subsequent behavioral preferences [30], and recent work has shown that larvae also exhibit appetite-dependent behavioral modifications [31]. We thus fed larvae ad libitum to fish food before testing their responses to each of the seven chemosensory cues (Fig 2B). Fed larvae showed no significant attraction to food (n=57, p=1), food extract (n=19, p=1), and RNA (n=20, p=1), supporting the prediction that microbial RNA functions as an attractant in the context of foraging. Fed larvae showed no defects in quinine-mediated aversion (n=24, p=0.003), demonstrating that the lack of response to foraging cues is not due to a global reduction in chemosensory behavior. Similar to starved larvae, fed animals showed no preference for the water control (n=39, p=1) or indole (100*μ*M n=36, p=0.87, 10mM n=17, p=1). Fed larvae exhibited significant aversion to o-cresol (n=36, p=0.024).

### Larval exploration behavior is best explained by a chemokinesis search model

Next we investigated the behavioral mechanism by which *Ae. aegypti* larvae locate sources of odor, since such information could provide insight into the chemosensory pathways that mediate the behaviors. We hypothesized that larval aggregation near attractive cues such as food is mediated by chemo-klino-taxis - a common form of directed motion observed in many animals and microbes [32, 33, 34]. In chemo-klinotaxis (hereafter chemotaxis), animals exhibit directed motion with respect to a chemical gradient. Alternatively, larvae may exhibit chemo-ortho-kinesis (hereafter chemokinesis) - a process in which animals respond to local conditions by regulating speed rather than direction - or chemo-klino-kinesis (hereafter klinokinesis) - in which animals respond to local conditions by regulating turning frequency. Finally, larvae may be unable to detect chemosensory stimuli, and thus exhibit purely random behavior (hereafter anosmic).To differentiate between these strategies, we quantified six observable metrics used to characterize navigation behavior. By identifying which variables correlate with stimulus preference, we can infer which search strategy best explains larval behavior (Table 1). Surprisingly, we found no evidence for chemotaxis near attractive or aversive chemicals. Starved larvae did not exhibit kinematic changes characteristic of chemotaxis, such as directional preference (ΔDP, p=0.18, Fig S3A). Further, larvae could not increase odor localization efficiency above random chance: discovery time for all cues was comparable across treatments (ΔD, p=1, Fig S3B). Larvae also did not perform klinokinesis: Turning frequency was unaffected by either the instantaneous concentration the larvae experienced (ΔCTI, p=1, Fig S3C) or change in concentration (ΔDTI, p=1, Fig S3D). Instead, we found that larval activity was most consistent with chemokinesis. Larvae altered movement speed when experiencing high local stimuli conditions (ΔCS, p<0.0001, Fig 3D) but not when moving up or down the concentration map (ΔDS, p=1, Fig S3E).

**Table 1:**
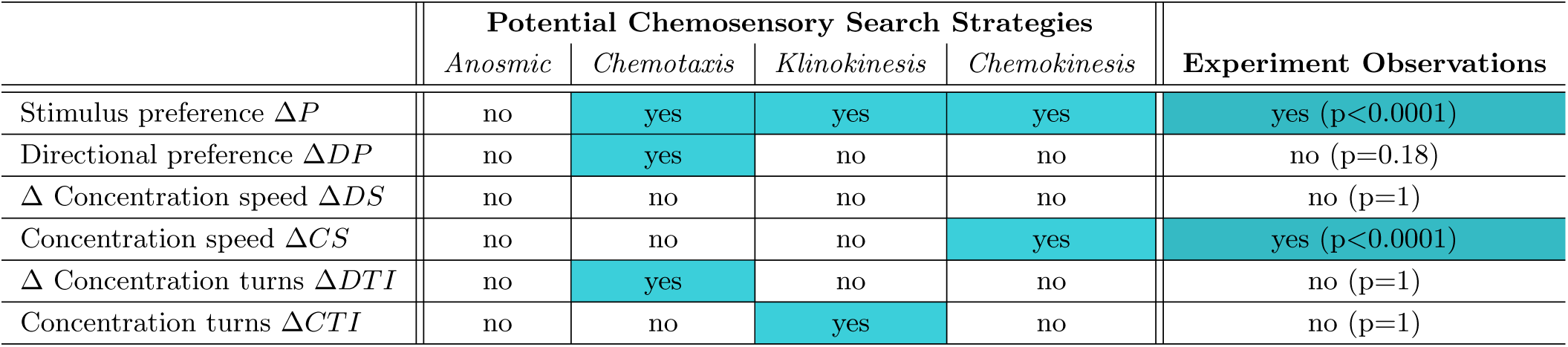
Comparing larval exploration behavior to canonical animal search strategy models. Four different chemosensory search strategies are listed (central columns) along with the expected observable behavior metrics for each strategy (left column). By comparing the experimental observations (right column) with the expected results, we determined that *Ae. aegypti* larval chemosensory navigation is best explained by an chemokinesis search strategy model.

**Figure 3:**
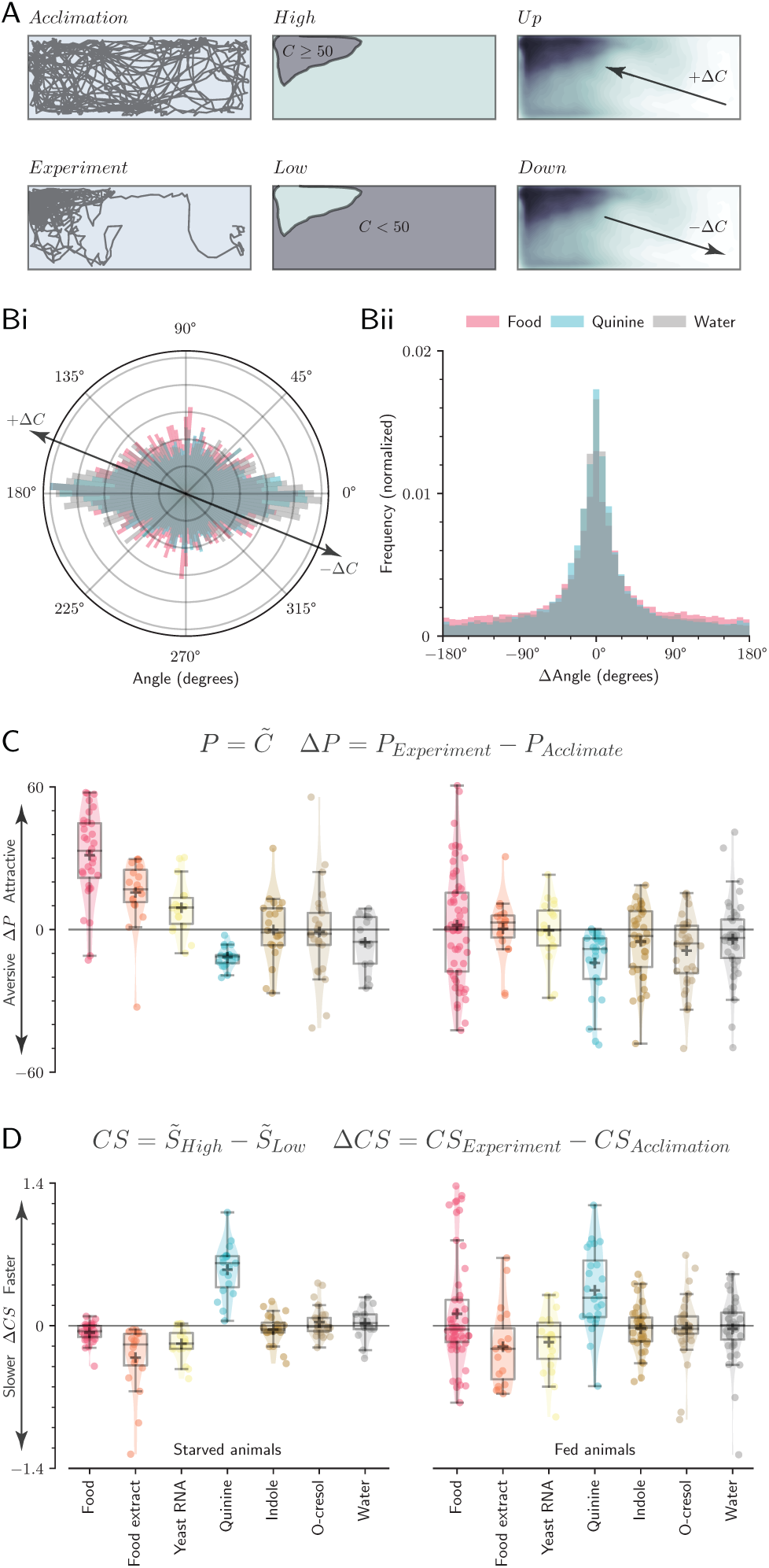
Larval exploration behavior is best explained by a chemokinesis search model. A: Diagram of behavioral quantifications. Larvae were observed during a 15-minute acclimation period in clean water, followed by a 15-minute experiment in the presence of the stimulus. The arena was divided into an area of high (≥50%) and low concentration (<50%). Larvae could move in a direction that increased local concentration (+ΔC) or decreased local concentration (-ΔC). Bi: Orientation of animals in the arena throughout the experiment. Larvae did not exhibit directional movement in response to appetitive or aversive stimuli. Note that larvae spend more time moving horizontally (0°, 180°) because the rectangular arena is longer in the horizontal direction. Bii: Larvae did not change frequency of turns (Δangle) in response to appetitive or aversive stimuli. C: Box plots for the population median (± 1 quartile), population mean (+ marker) and mean response for each individual (dots) for larval preference (ΔP). A horizontal line at 0 represents no change in behavior following stimulus addition. D: As in C, except for stimulus-dependent changes in Concentration-dependent Speed (ΔCS).

### Chemokinesis is superior to chemotaxis for avoiding repellents in realistic larval environments

Our results were particularly surprising considering that many insects use chemotaxis rather than chemokinesis to navigate [35]. Could chemokinesis be unusually advantageous in microhabitats, such as those utilized by mosquitoes in urban environments? We developed four data-driven models to simulate larval activity using chemokinesis, klinokinesis, an anosmic random walk, or chemotaxis. In these data-driven models, larval speed and turn angle was determined at each time step from a bootstrap resampling of empirical data from all larval trajectories in clean water (n=248 larvae during the acclimation phase, fed ad libitum. n=445,925 trajectory data points). This extensive empirical dataset allowed us to investigate the success of each search strategy while retaining the characteristics of authentic larval behavior. In addition, we created an exponential regression model to simulate diffusion properties observed in our experimental arena (p<0.0001, Fig S2E). Using these data-driven representations of larval speed, larval turning rate, and chemical diffusion in naturalistic larval habitats, we compared the success of each simulated search strategy in two separate challenges: a foraging task measuring time elapsed before finding a food source, and a repellent-avoidance task measuring the proportion of time spent in high-repellent environments. The success of each search strategy was explored across a range of common habitat sizes observed in urban environments [36] (Table 2). If larval chemokinesis is an adaptation to small urban microhabitats, we expect the chemokinesis search model to perform better than other strategies, and for this difference to be more apparent at smaller habitat sizes. Indeed, we found that chemokinesis was by far the best strategy in the repellent-avoidance task when avoiding potentially stressful environments (e.g. toxins, pollutants, or pesticides). Further, the difference between strategies was greatest at small habitat sizes (Fig 5A,B).

**Figure 4:**
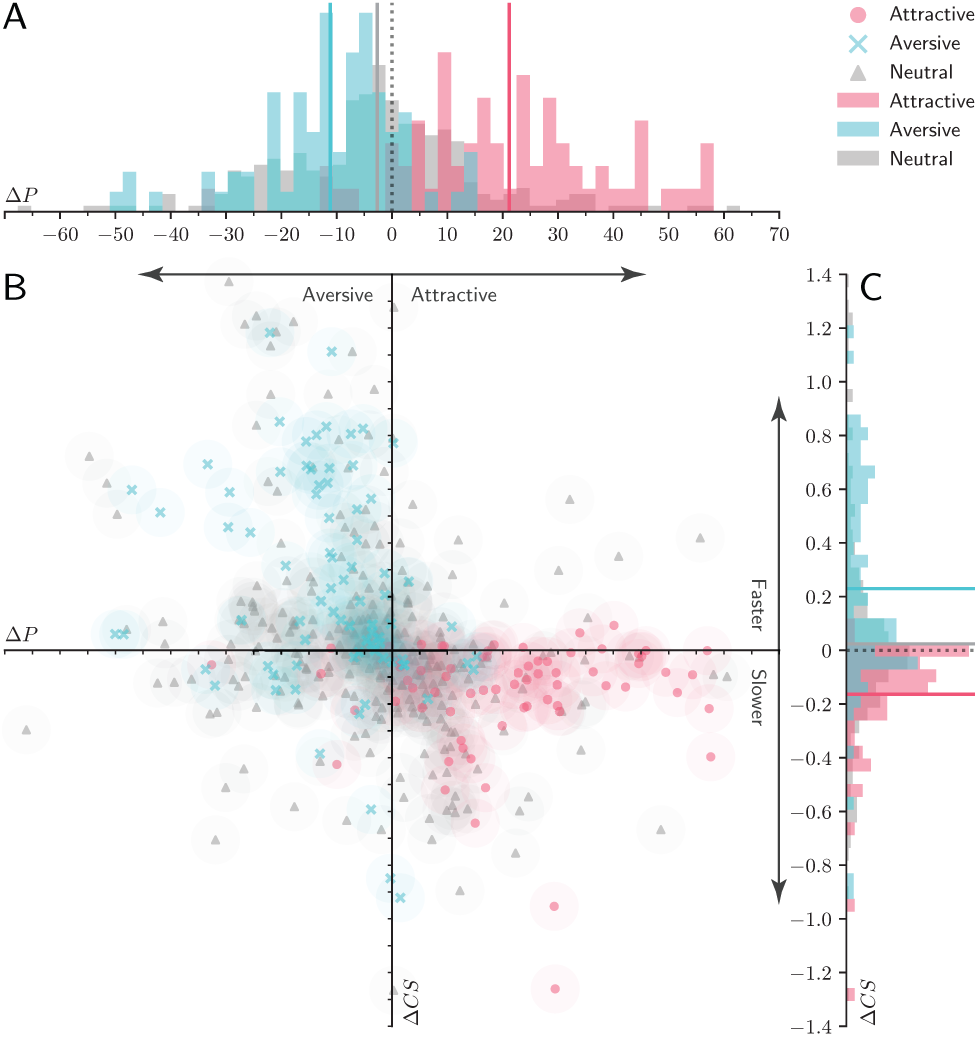
Larval stimulus preference is correlated to concentration-dependent movement speed. A: Larval preference (ΔP) significantly correlates with Concentration-dependent Speed (ΔCS). Results from all experiments are shown grouped into three categories: attractive (pink: food, food extract, and yeast RNA in starved larvae), aversive (blue: quinine), and neutral (grey: water, indole, o-cresol in fed and starved larvae; food, food extract, and yeast RNA in fed larvae). B: Normalized frequency histograms of ΔP. Mean response to aversive, neutral, and appetitive cues are visualized as solid vertical lines in the corresponding color. A dotted black line at zero indicates the expected outcome if the added stimulus had no effect on larval behavior. C: As in B, except for normalized frequency histograms of larval ΔCS.

**Table 2:** Ecologically realistic habitat sizes analyzed through computational modeling. A range of habitat sizes were selected from a literature search of realistic habitat sizes for *Ae. aegypti* larvae ([36] and references therein).

|  | Radius | Frequency | Examples |
| --- | --- | --- | --- |
| i | <5cm | 27.8% of habitats | Ant traps |
| ii | 5-9cm | 9.7% of habitats | Tin cans, bottles |
| iii | 9-17cm | 32.3% of habitats | Jars, bowls, vases |
| iv | 17-20cm | 3.1% of habitats | Plates, pails |

### Starved Aedes aegypti optimize exploration behavior to increase the probability of finding food

In contrast, chemokinesis was also the worst strategy in the foraging task, taking over an hour to find the simulated food source (Fig 5C,D). However, our data-driven models resampled empirical data collected from animals fed ad libitum. Many organisms change their speed or activity rate when starved [37], and we predicted that starved *Ae. aegypti* may also alter their exploration behavior to increase the probability of discovering food [37]. Experimental observations showed evidence for starvation-mediated behavior changes - starved animals spent more time exploring (p<0.0001, Fig 6A) and spent less time near walls and corners (p<0.0001, Fig 6B). If these starvation-mediated behavioral changes are adaptative, we expect the data-driven chemokinesis model to perform much better at the foraging task when given empirical data from starved larvae. Thus we tested the success of each search model in the foraging task using bootstrap resampling of empirical data from starved animals (n=168 starved larvae during the acclimation phase, n=302,096 trajectory data points). The starved chemokinesis model discovered the food source almost an hour faster across all habitat sizes (Fig 6C), supporting our hypothesis that starvation-mediated changes in larval behavior increase the probability of finding food.

**Figure 5:**
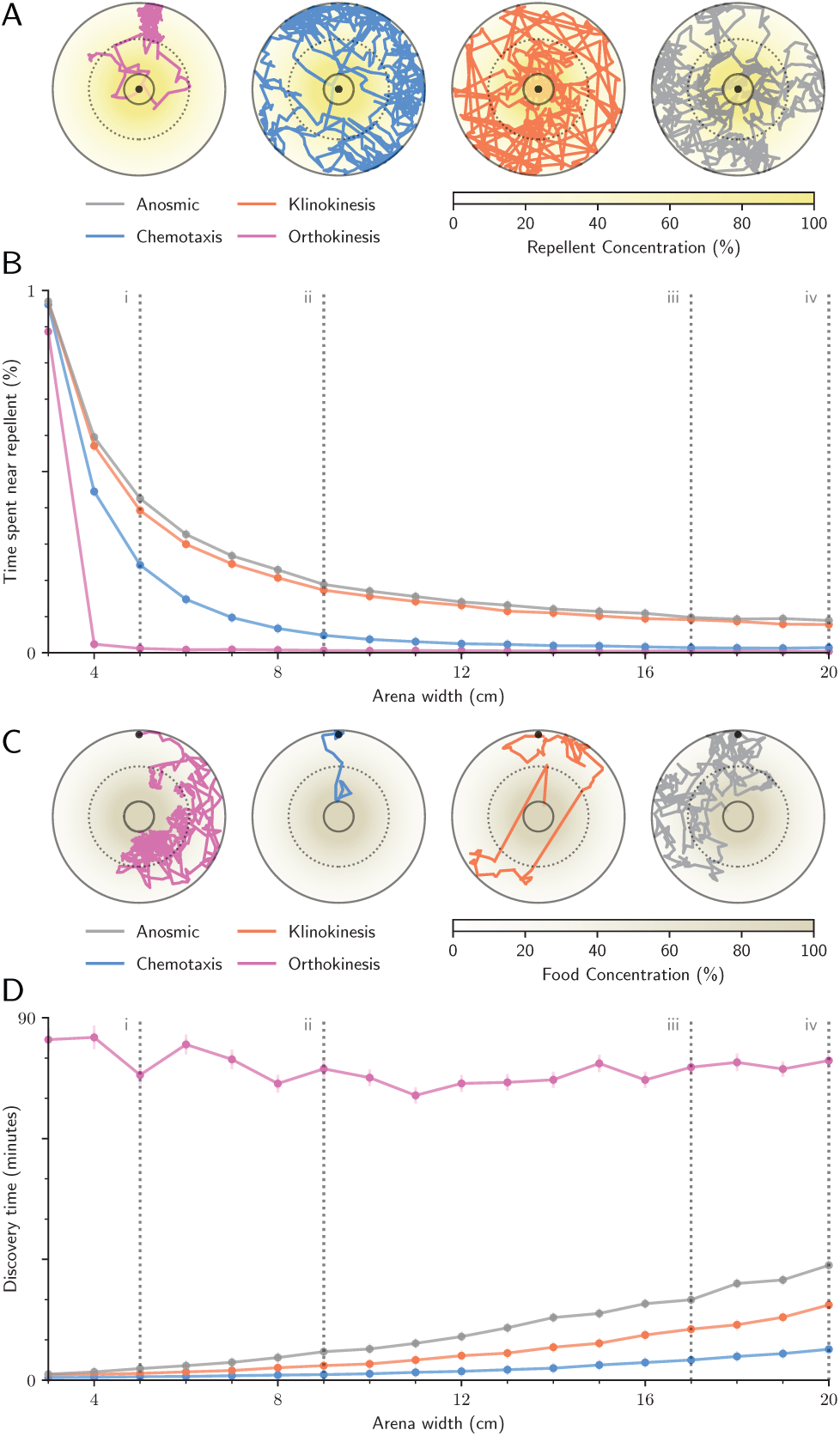
Chemokinesis is superior to chemotaxis for avoiding repellents in realistic larval environments. A: Sample trajectories for the repellent-avoidance task. B: Success of each search strategy in the repellent-avoidance task (mean ± standard error). C: Sample trajectories for the foraging task. (A,C): Dotted lines mark 50% concentration. Foraging trajectories begin at the top of the 6cm-diameter arena, and repellent-avoidance task trajectories at the arena center (black dot). Starting point was randomized in actual analyses. D: Success of simulated search strategies in the foraging task. (B,D): Dashed grey lines correspond to ecologically relevant habitat sizes described in Table 2.

**Figure 6:**
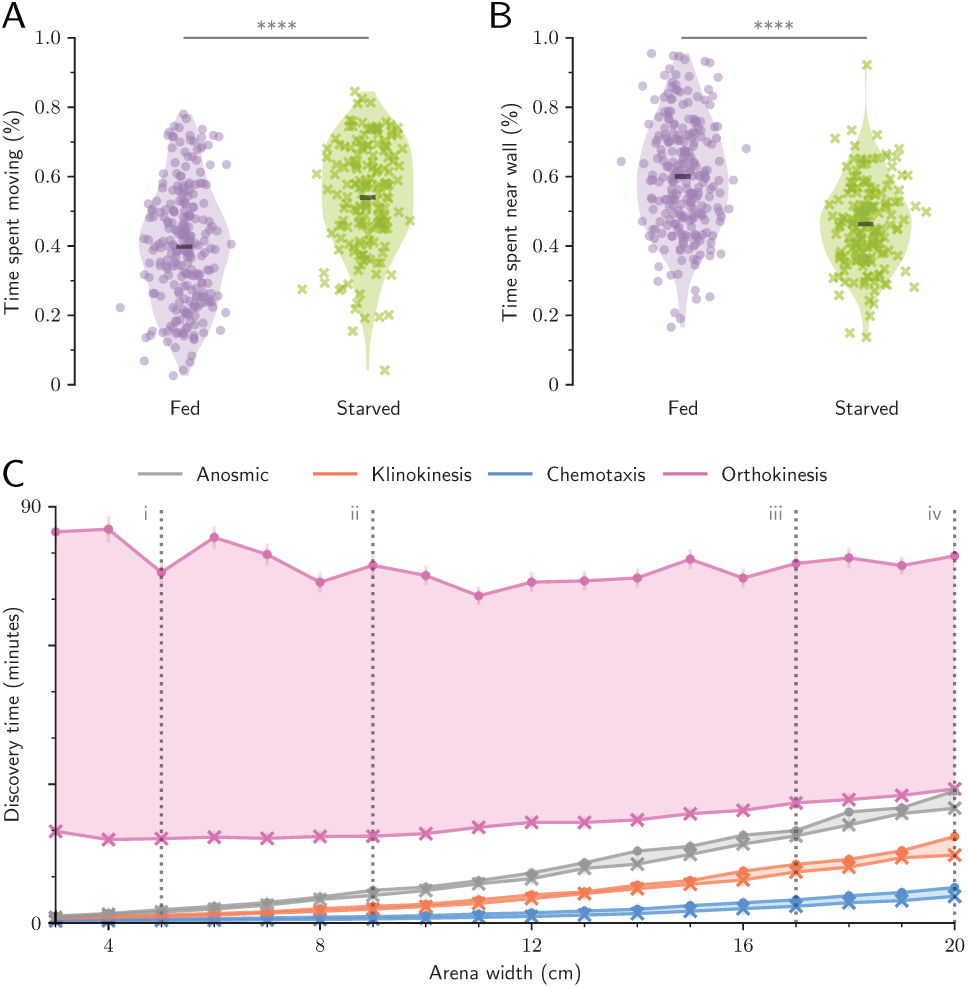
Starved *Ae. aegypti* optimize exploration behavior to increase the probability of finding food. A: Starved larvae spend significantly more time exploring the arena than fed larvae. B: Starved larvae spend significantly less time within one body length of an arena wall. (A,B) Violin plot. Dots are the means for each individual, and black bar is the mean across all individuals (n>168 per treatment); asterisks denote p<0.0001 (Welch’s t-test). C: Simulated chemokinetic larvae using empirical data from starved animals found the food source consistently faster than the same model using data from fed animals. Shaded regions show difference between fed (X markers) and starved (dots) simulations (mean ± standard error). Dashed grey lines correspond to ecologically relevant habitat sizes described in Table 2.

Nevertheless, starved chemokinesis simulations still performed worse than all other strategies in the foraging task. This result, coupled with the runaway success of the chemokinesis model in the repellent-avoidance task, suggests that avoiding repellents may be particularly important for *Ae. aegypti* larval fitness. If avoiding repellents is essential for *Ae. aegypti*, any starvation-mediated behavioral adaptations may be constrained by the additional requirement of retaining successful repellent-avoidance behavior. If so, we would expect to see very little difference in repellent-avoidance success across simulations based on empirical data from fed or starved larvae. Our results supported these predictions: Although starved simulations performed slightly worse compared to fed simulations, the difference was small: starved simulations only spent an average of 1% more time near the repellent (Fig S4C).

## Discussion

In this study we quantify essential characteristics of *Ae. aegypti* larval behavior that are crucial for the development of future studies. Further, we identify previously unknown behaviors that highlight the unique evolutionary history and developmental biology of these disease vector mosquitoes. First, we show that larvae perceive microbial RNA as a foraging attractant, but do not respond to several olfactory cues that attract adult *Ae. aegypti* for oviposition. Second, we demonstrate that *Ae. aegypti* larvae use chemokinesis, rather than chemotaxis, to navigate with respect to chemical sources. Using data-driven computational modeling, we further show that chemokinesis is superior to chemotaxis in avoiding repellents in ecologically relevant larval habitat sizes. Finally, we use experimental observations and computational analyses to demonstrate that larvae respond to starvation pressure by changing their behavior to increase the probability of finding a scarce food source without compromising their ability to successfully avoid repellents.

These results are fascinating from both a developmental biology and disease prevention perspective. In their adult form and during flight, *Ae. aegypti* exhibit an odor-tracking behavior termed odor-conditioned optomotor anemotaxis, where encounter with an odor gates an upwind surge in the wind direction [38]. In this behavior, successive odor encounters are necessary to prolong the upwind flight towards the upwind odor source, and the gradient information is not necessary to elicit the upwind responses. In other insects, such as *D. melanogaster*, while walking but not flying, these animals exhibit a form of chemotactic behavior where bilateral comparisons are made between antenna [39]. It remains unclear whether walking adult *Ae. aegypti* may also exhibit similar chemotactic behaviors, but given the differences between adult and larval responses, this species may provide an excellent developmental model to identify neurobiological pathways integral to olfactory navigation. Previous studies on mosquito larvae can further contextualize our results and provide additional insight. Unlike *Ae. aegypti, Anopheles gambiae* mosquito larvae prefer both indole and o-cresol, in addition to many other olfactory stimuli [23]. The stark differences in larval chemosensory behavior mirror the many differences observed between the adults of these two species [40], and suggests that studies should be cautious of generalizing among disease vector mosquitoes.

Although adult *Ae. aegypti* feeding is regulated by ATP perception [41], we are unaware of other work demonstrating RNA attraction in *Ae. aegypti* larvae. In our state-dependent preference experiments, we investigate the ecological basis of larval RNA attraction, and propose that RNA may function as a foraging indicator in the larval environment. Although the receptor responsible for RNA detection is unknown, work in *D. melanogaster* suggests that a gustatory or ionotropic receptor may be more likely candidates than an olfactory receptor. In addition, an earlier study demonstrated that olfactory deficient (*orco -/-) Ae. aegypti* larvae showed no defects in attraction to food or avoidance of quinine [22]. Taken together, our results support the hypothesis that sensory information gained from gustatory or ionotropic receptors may be more integral to larval chemosensation than olfactory receptors. Further, larval attraction to RNA suggests that the importance of nucleotide phagostimulation is preserved throughout a mosquito’s life cycle, from larval foraging to adult blood engorgement and oviposition.

Our computational experiments suggest an ecological basis for the lack of chemotaxis in *Ae. aegypti* larvae. Although our experiments showed that chemotaxis is superior to chemokinesis in foraging, chemokinesis surpassed chemotaxis, klinokinesis and anosmic strategies in avoiding repellents. This suggests that the role of chemosensation in larvae is primarily tuned toward aversive responses. Indeed, known characteristics of larval physiology support this idea. Although larvae can survive for up to a week without food, they quickly succumb to toxic bacterial byproducts [10, 42]. We propose that *Ae. aegypti* larvae combat starvation pressure primarily through physiological adaptations such as fat stores, and resist toxins in the environment through chemosensory behaviors optimized for avoiding repellents.

Our study also raises a number of comparative questions that could be addressed in future research. For instance, is chemokinesis in mosquito larvae associated with generalized spatial restriction, or with human association and man-made containers in particular? Future studies could compare chemotactic ability in other spatially constrained mosquitoes, such as *Toxorhynchites* (which inhabit tree holes) or *Aedes albopictus* (another container-breeding mosquito) [43], to species that oviposit in larger bodies of water such as *Aedes togoi* (marine rock pools) or opportunistic species such as *Culex nigripalpus* that oviposit in a wide range of habitat sizes [43, 44]. Additionally, computational modeling of fluid dynamics and larval movement may help determine whether chemotaxis is physically challenging in small, man-made environments. Shallow gradients in small containers may diffuse too quickly to be used as a reliable chemical signal - particularly considering our results showing that larval movement significantly increases stimulus diffusion [45].

Synanthropic mosquitoes are increasingly important to global health as urbanization progresses: Currently over half of all humans live in urban environments, and this proportion is only expected to increase [46]. Adaptations that facilitate human cohabitation, like specialized larval foraging strategies, are vital to our understanding of mosquito behavior and success as a disease vector [9].

## Materials and Methods

### Insects

Wild-type *Ae. aegypti* (Costa Rica strain MRA-726, MR4, ATCC Manassas Virginia) were maintained in a laboratory colony as previously described [47]. Experiment larvae were separated within 24 hours of hatching and reared at a density of 75 per tray (26×35×4cm). One day before the experiment, 4-day-old larvae were isolated in Falcon*™* 50mL conical centrifuge tubes (Thermo Fischer Scientific, Waltham, MA, USA) containing ~15mL milliQ water. Starved larvae were denied food for at least 24 hours before the experiment. Animals that died before eclosion or pupated during the experiment were omitted. Because it was not possible to detect younger larvae using our video tracking paradigm, we mitigated possible age-related behavioral confounds by standardizing the age of experimental larvae.

### Behavior arena and experiment

We previously developed a paradigm to investigate chemosensory preference in larval *Ae. aegypti* [22]. In this study we expanded our protocol by mapping the chemosensory environment in our arena using fluorescein dye. 100μL of fluorescein dye was added to a white arena of the same material and dimensions, each containing one *Ae. aegypti* larva. Dye color was converted to concentration values using a standardization dataset of 13 reference concentrations (Fig S2C). Dye diffusion through time was quantified by the mean of all values in each 1mm^2^ area, linearly interpolated throughout time (n=10, Fig S2B). During behavior experiments, we recorded animals for 15 minutes before each experiment to analyze baseline activity and confirm that the arena was fair in the absence of chemosensory cues. Subsequently, 100pL of a chemical stimulus was gently pipetted into the left side of the arena to minimize mechanosensory disturbances, and larval activity was recorded for another 15 minutes (Fig 1C).

### Selection and preparation of odorants

Odorants (indole, o-cresol) were prepared at 100μM in milliQ water (Aldrich #W259306; Aldrich #44-2361). Indole was also prepared similarly at 10mM. Quinine hydrochloride was prepared at 10mM in milliQ water (Aldrich #Q1125). Larval food (Petco; Hikari Tropic First Bites) was prepared at 0.5% by weight in milliQ water and mixed thoroughly before each experiment to resuspend food particles. To prepare the food extract solution, 0.5% food was dissolved in milliQ water for one hour and filtered through a 0.2*μ*m filter (VWR International #28145-477). For the yeast RNA solution, total RNA from *Saccharomyces cerevisiae* yeast was prepared at 0.1% by weight in DEPC-treated, autoclaved 0.2*μ*m filtered water (Aldrich #10109223001; Ambion #AM9916). Yeast RNA, food, and food extract were prepared fresh daily. Although chemicals diffuse at different rates depending on molecular size and physico-chemical properties, diffusion coefficients in water were unavailable for the majority of chemicals tested. Therefore, it is important to note that our chemical diffusion map is an approximation of the actual chemosensory environment experienced by larvae.

### Video Analyses

Video data was obtained and processed as previously described [22] using Multitracker software by Floris van Breugel [48] and Python version 3.6.2. Additionally, approximate larval length was measured for each animal in ImageJ Fiji [49], as the pixel length from head to tail, in a selected video frame that showed the larva in a horizontal position. Lengths were converted to mm using the known inner container width as the conversion ratio. Experimenters were blind to larval sex when measuring lengths.Throughout our analyses, the arena was divided into areas of high concentration (≥50% initial stimulus) and low concentration (<50%). Larvae could move in a direction that increased local concentration or decreased local concentration. We discounted concentration changes caused by diffusion while the larvae remained immobile. A threshold of Δ2%/s was required to qualify as moving up or down the concentration map.

### Statistical Analyses

Statistical analyses were performed in R version 3.5.1 [50]. A Bonferroni-Holm correction was applied to all statistical analyses. A Mann-Whitney test was used to compare body length of fed and starved males and females (Fig S1A). Linear least squares regression was used to assess the effect of time of day to animal speed, time spent moving, and time spent near walls during the acclimation phase (Fig S1B-D). Paired-samples Welch’s t-tests were used to compare the median chemical concentration chosen by the larvae throughout the 15-minute experiment to the behavior of the same larvae throughout the 15-minute acclimation phase. This preference metric was also quantified a single value (ΔP, P_*Experiment*_-P_*Acclimation*_, Fig 3, Fig 4). For all subsequent analyses on behavioral mechanisms, larval behavior during the acclimation phase was subtracted from larval activity during the experiment phase to normalize for differences between individuals and larval preference for corners and walls. When investigating potential differences between attraction and aversion behaviors, we grouped stimuli into cues that elicited significant attraction (ΔP>0, p<0.05), significant repulsion (ΔP<0, p<0.05), or neutral response (p≥0.05). A Kruskal-Wallis test was used to compare behavioral metrics among these three stimuli classes (Fig 3D, Fig 4, Fig S3). These other behavioral metrics included directional Preference (ΔDP), defined as the difference in time moving up or down the concentration map; Discovery time (ΔD), defined as the time elapsed before initial encounter of high (≥50%) concentration of the stimulus; Concentration-dependent Speed (CS), defined as the difference in speed at high (≥50%) and low (<50%) local concentrations; ΔConcentration-dependent Speed (ΔDS), defined as the difference in speed while moving up or down the concentration map; Concentration-dependent Turn Incidence (ΔCTI), defined as the difference in turning rate (turns per second, turns defined as instantaneous change in angle of >30°) at high and low local concentrations; and ΔConcentration-dependent Turn Incidence (ΔDTI), defined as the difference in turning rate while moving up or down the concentration map. For statistical analyses, larvae that never entered areas of high concentration were assigned a ΔD of 15 minutes, corresponding to the end of the experiment, and a ΔCS and ΔCTI of 0 (placeholder values chosen to reduce Type I error).

### Computational Modeling

We developed four data-driven models to investigate larval exploration success in different environments. The empirical dataset used in these models represented all data points taken from larvae observed in clean water before the addition of experimental stimuli (n=248 fed, n=168 starved). In the foraging task, simulated animals explored until they encountered a food source at the center of the arena (scaled to arena size, comprising 3% of total area). This discovery time was recorded for each of 1000 simulations per arena size and per model. In the repellent-avoidance task, simulated larvae explored for 15 minutes, and the percentage of time spent within ≥50% of the repellent was recorded. We defined the simulated chemical boundary conditions using an exponential regression model of distance and concentration based on our chemical map data (Fig S2E). All simulated larvae began at a random point within the arena. In the anosmic model, instantaneous speed and angle was randomly sampled from the empirical dataset and applied to the larval trajectory at each time step (2fps). The chemokinesis model explored while sampling chemical concentration. In this model the empirical dataset of instantaneous speed was sorted and split into slow and fast halves. If food concentration was ≥50% (or repellent concentration was <50%), speed was sampled from the slow half. If food concentration was <50% (or repellent concentration was ≥50%) speed was sampled from the fast half. In the chemotaxis model, if food concentration increased by ≥1% (or repellent concentration decreased by ≥1%), the animal continued in the same direction for the next movement step. Similarly, for klinokinesis the animal continued in the same direction for the next movement step if the local concentration was ≥50% (foraging task) or <50% (repellent-avoidance task). For chemotaxis we simulated a range of biologically plausible concentration sensitivities ranging from 0.1% to 10% and found that this did not affect our conclusions (Fig S4A,B).

## Acknowledgements

This work was supported in part by the National Institute of Health grant 1RO1DCO13693-04 to J.A.R.; National Science Foundation grants IOS-1354159 to J.A.R. and DGE-1256082 to E.K.L.; Air Force Office of Sponsored Research under grant FA9550-16-1-0167 to J.A.R.; and the Robin Mariko Harris Award to E.K.L. We thank Floris van Breugel for assistance with video data analysis, the University of Washington Biostatistics Consulting Group for statistical advice, and Binh Nguyen and Kara Kiyokawa for maintaining the Riffell lab mosquito colony. We also thank Thomas Daniel, Bingni Brunton, Kameron Harris, and the Kincaid 320 Python Club for insightful discussions on programming and data management.

## Author Contributions

Conceptualization: E.K.L. and J.A.R.; Methodology: E.K.L. and J.A.R.; Software: E.K.L.; Investigation: E.K.L.and T.S.G.; Resources: E.K.L. and J.A.R.; Data Curation: E.K.L; Writing - Original Draft: E.K.L; Writing-Review Editing: E.K.L, J.A.R, and T.S.G.; Visualization: E.K.L; Supervision: J.A.R; Project administration: J.A.R; Funding acquisition: E.K.L. and J.A.R.

## Supplementary materials

### Supplementary Data and Code

All code is available for download at github.com/eleanorlutz/aedes-aegypti-2019

**Figure S1:**
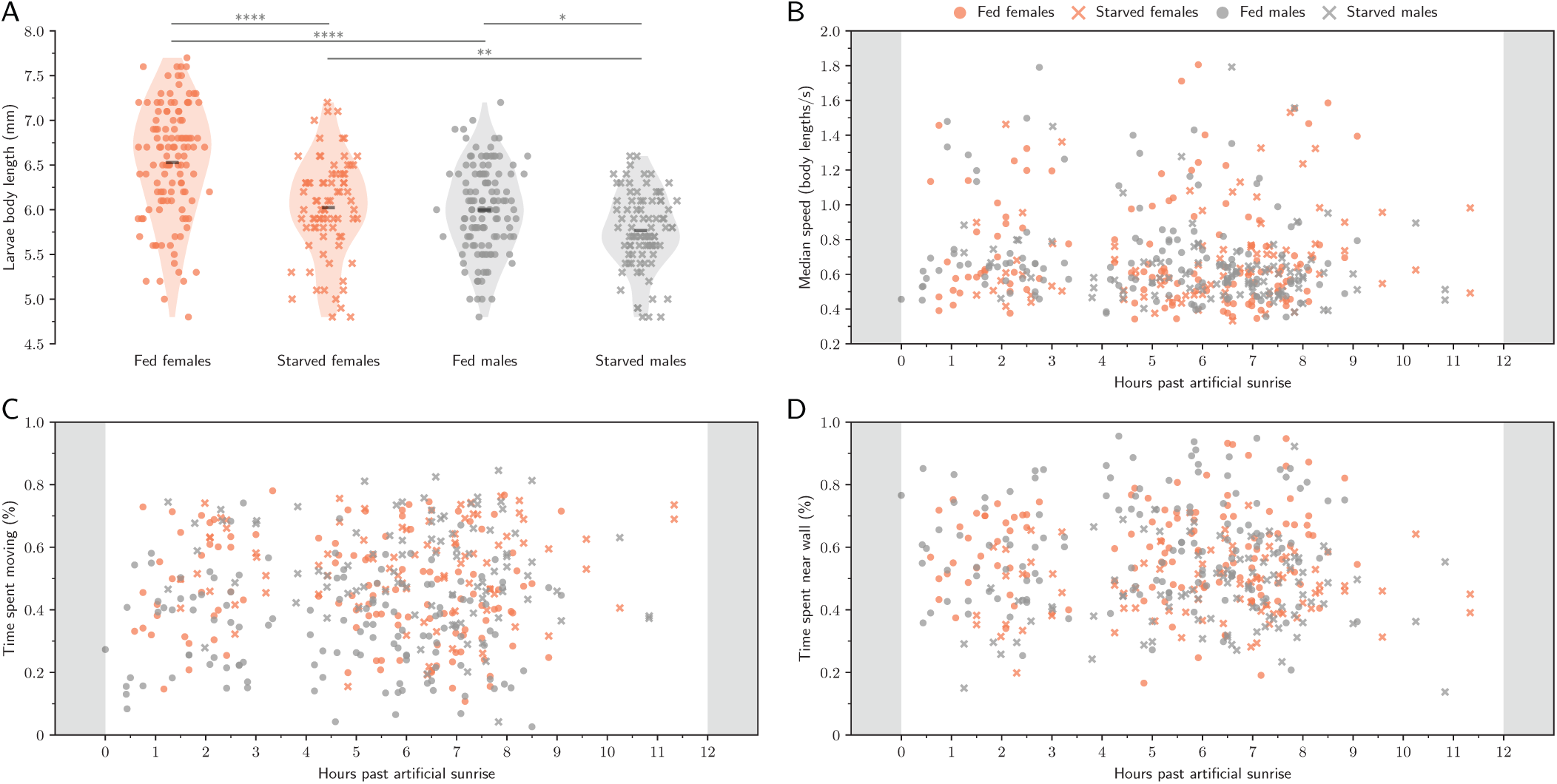
Effects of sex, physiological state, and circadian timing on larval physiology. A-D: Fed females (orange dots, n=120) and males (grey dots, n=128), starved females (orange X markers, n=79) and males (grey X markers, n=89). A: Violin plot. Scatter points show the body length (mm) for each individual, and the black bar is the mean across all individuals; asterisks denote significance values (Welch’s t-test). Larval body length is influenced by sex and starvation state. B: No change was observed in median speed (body lengths/s). Note that the sampling rate throughout the day was not consistent due to the work schedule of experimenters involved in the project. C: No change was observed in time spent moving throughout daylight hours. D: No change was observed in proportion of time spent within one body length of the wall throughout daylight hours.

**Figure S2:**
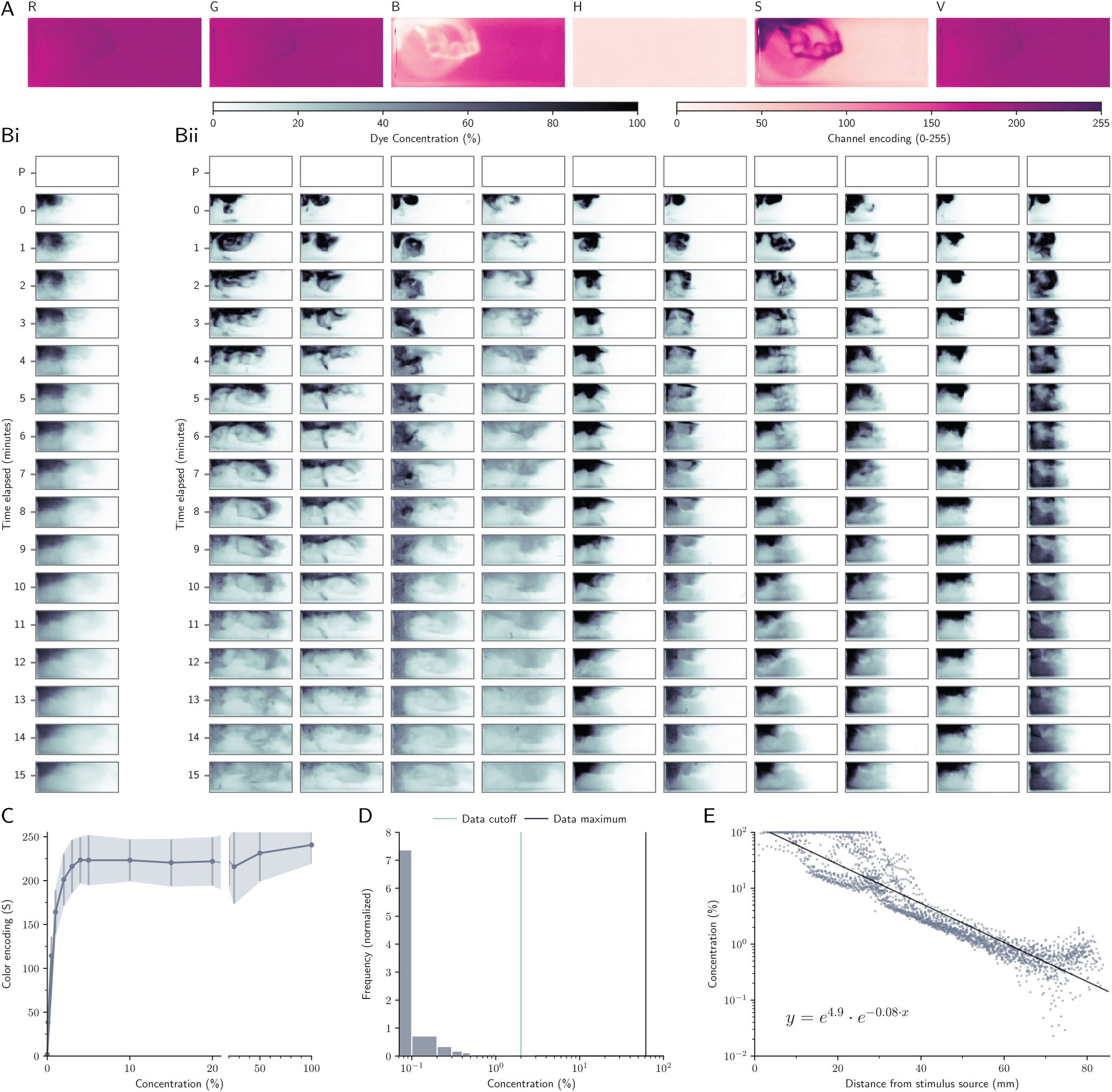
Creating a concentration gradient map to analyze and model larval search behavior. A: To quantify fluorescein dye diffusion, photographs were taken every minute using a Canon PowerShot ELPH 320 HS camera. Of the available color information channels (RGB, HSV), the saturation channel (S) contained the most information and was used to represent dye color throughout image analyses. Bi: Dye diffusion through time was quantified by the mean of all values in each 1mm2 area, linearly interpolated through time (n=10 experiments containing larvae). A control photograph was taken before the start of each experiment (P) but was not used to construct the chemical gradient map. Bii: Individual variation between trials. Each column represents data from one experiment through time. C: Dye color (S) was converted to raw concentration values using a standardization dataset of 13 reference concentrations. 20mL of each reference concentration was poured into the entire arena and photographed. D: Because 100*μ*L of dye is immediately diluted in the 20mL behavior arena water volume, reference concentration colors could not be used to directly convert color to % maximum concentration. Instead, the maximum concentration value was normalized to ≥95% of all color measurements across all experiments. E: To create a concentration map for computational simulations in different arena sizes, we analyzed the relationship between concentration and distance from stimulus source at time=0. Concentration values for individual 1×1mm^2^ sections across all 10 experiments at time=0 (dots).

**Figure S3:**
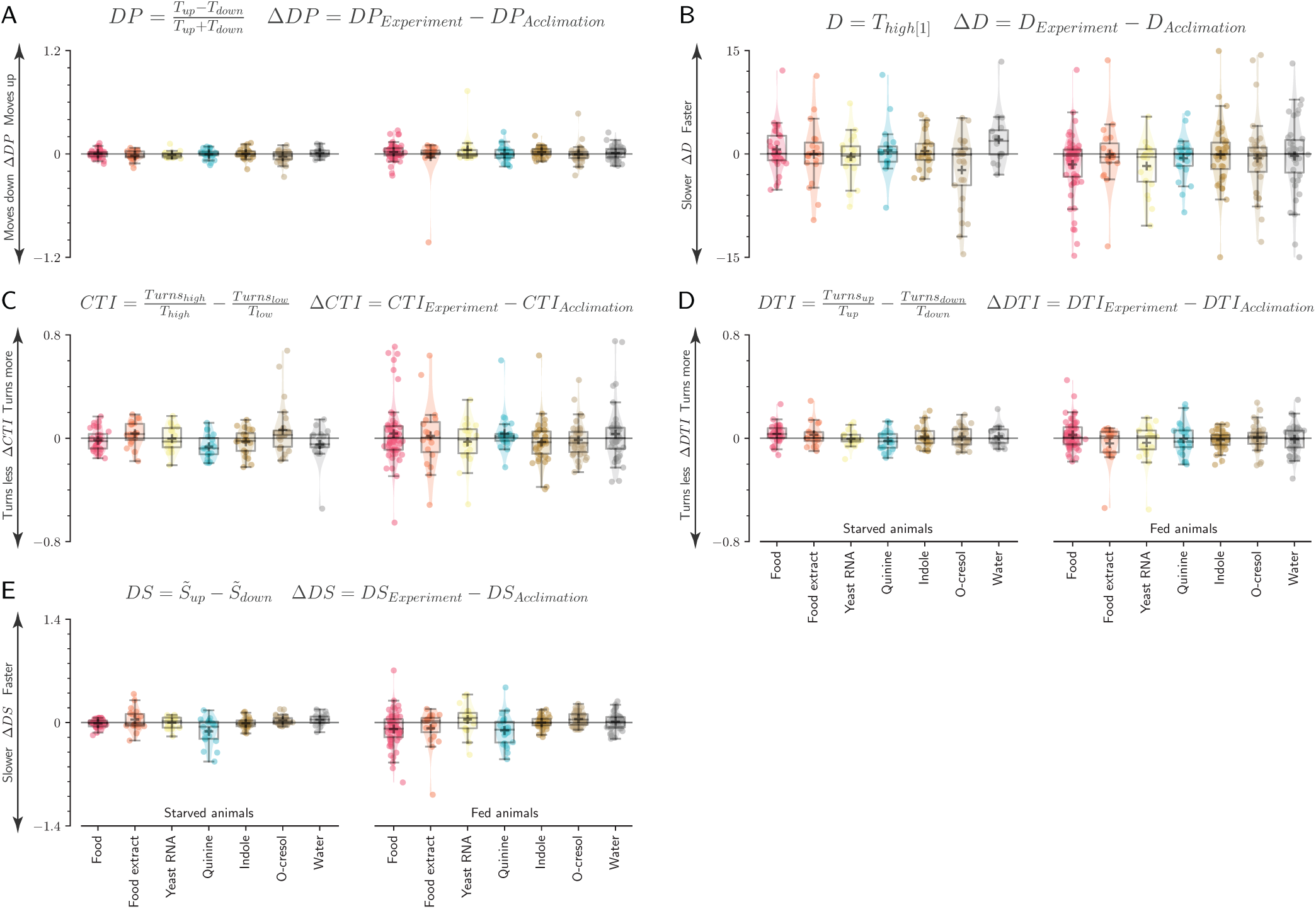
Larval behavior is not consistent with chemotaxis or klinokinesis search strategy models. A-E: Box plots for the population median (± quartile), population mean (+ marker) and mean response for each individual (dots). We observed no significant changes across stimuli for any of these five behavioral metrics (p>0.05, Kruskal-Wallis test). A: Directional Preference ∆DP, difference in time (*T*) moving up or down the concentration map. B: Discovery time ∆D, time (*T*) elapsed before initial encounter of high concentration (≥50%). C: Concentration-dependent Turn Incidence ∆CTI, difference in turning rate at high and low local concentrations. D: ∆Concentration-dependent Turn Incidence ∆DTI, difference in turning rate while moving up or down concentration.E: ∆Concentration-dependent Speed ∆DS, difference in mean speed 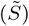 while moving up or down the concentration map.

**Figure S4:**
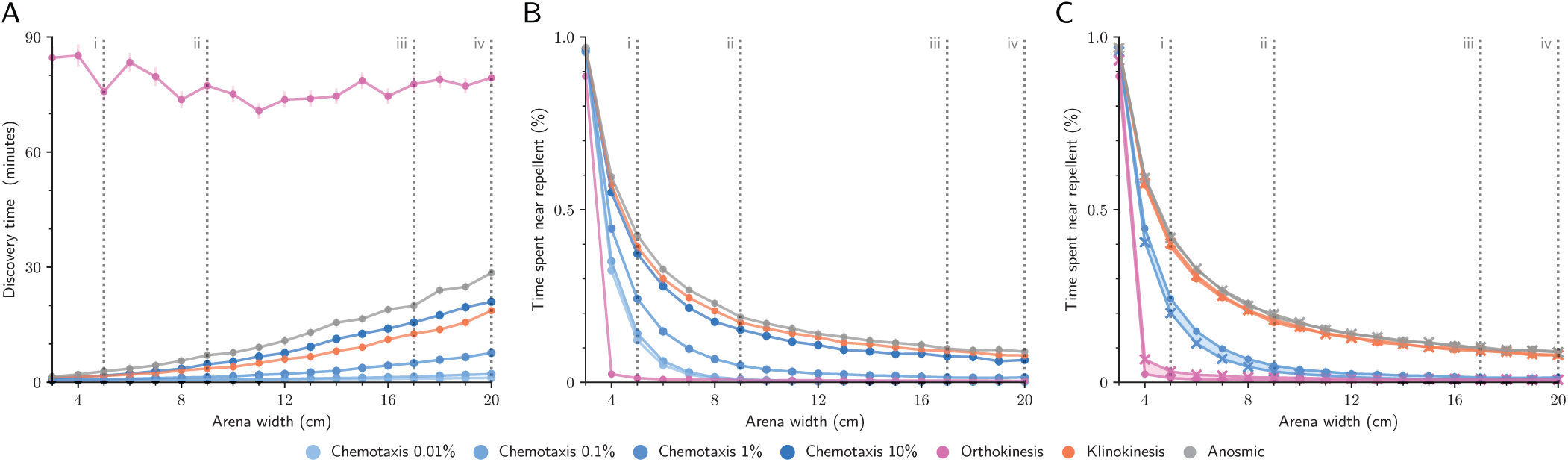
Simulation results are not affected by chemotactic sensitivity, or by substituting the starved and fed empirical datasets in the repellent-avoidance task. A: Time elapsed before simulated larvae discovered food in the foraging task (mean ± standard error). Chemotaxis % values indicate the lowest concentration difference detectable by simulated larvae during each time step (2fps). B: Time spent in high-repellent areas during the repellent-avoidance task (mean ± standard error). All chemotactic sensitivities performed worse than the chemokinesis model. C: Starved simulations (X markers) and fed simulations (dots) performed similarly well during the repellent-avoidance task (mean ± standard error, shaded regions show difference between fed and starved simulations). In all panels, dashed grey lines correspond to ecologically relevant habitat sizes described in Table 2.

